# Neural oscillations while remembering traumatic memories in PTSD

**DOI:** 10.1101/639310

**Authors:** Inbal Reuveni, Noa Herz, Omer Bonne, Tuvia Peri, Shaul Schreiber, Yuval Harpaz, Abraham Goldstein

**Author notes:** contributed equally (Inbal Reuveni, Noa Herz). Corresponding author: Abraham Goldstein, Gonda Brain Research Center, Bar-Ilan University, Ramat Gan, 52900, Israel.

## Abstract

**Background:** In posttraumatic stress disorder (PTSD), the traumatic event is often re-experienced through vivid sensory fragments of the traumatic experience. Though the sensory phenomenology of traumatic memories is well established, neural indications for this qualitative experience are lacking. The current study aimed at monitoring the oscillatory brain activity of PTSD patients during directed and imaginal exposure to the traumatic memory using magnetoencephalography (MEG), in a paradigm resembling exposure therapy.

**Methods:** Brain activity of healthy trauma-exposed controls and PTSD participants was measured with MEG as they listened to individualized trauma narratives as well as to a neutral narrative and as they imagined the narrative in detail. Source localization analysis on varied frequency bands was conducted in order to map neural generators of altered oscillatory activity.

**Results:** PTSD patients exhibited increased power of high-frequency bands over visual areas and increased delta and theta power over auditory areas in response to trauma recollection compared to neutral recollection, while controls did not show such differential activation. PTSD participants also showed abnormal modulation of lower frequencies in the medial prefrontal cortex.

**Conclusions:** Elicitation of traumatic memories results in a distinct neural pattern in PTSD patients compared to healthy trauma-exposed individuals. Investigating the oscillatory neural dynamics of PTSD patients can help us better understand the processes underlying trauma re-experiencing.

## Introduction

The nature and underlying mechanism of traumatic memories has been a matter of controversy for over a century. The effect of traumatic events on the neural system was already noted by James, who stated that: “an impression may be so exciting emotionally as almost to leave a scar upon the cerebral tissues” (James, 1890). In post-traumatic stress disorder (PTSD), a traumatic experience leads to the development of intrusive memories, described as qualitatively different from other ordinary or stressful memories. These memories tend to be fragmentary, and consist of isolated visual, auditory, olfactory, or tactile sensations, with visual intrusions being the most common type of sensory intrusions across all types of trauma (Ehlers et al., 2002). These memories are exceptionally vivid and lack time perspective, and may therefore be experienced as something that is happening in the present rather than in the past (Ehlers et al., 2004). Clinical accounts consistently include such emotional and perceptual elements as their prominent features over and above declarative components. Based on these observations a dual representation theory has been proposed (Brewin et al., 2010), depicting two parallel memory systems: verbally accessible memory (VAM) and situationally accessible memory (SAM). The VAM system supports declarative representations of the event within its associated autobiographical context in a form accessible to deliberate retrieval and manipulation. The SAM, on the other hand, contains detailed sensory and perceptual images that can be accessed only involuntarily and are the source of intrusions conveying the traumatic moments. According to this model, traumatic events are stored as SAMs, without the association to the VAM system that is usually found in non-traumatic events. This allows retrieval of SAMs triggered by environmental or internal cues reminiscent of the trauma without retrieval of the appropriate autobiographical context (4). Another physiological model for re-experiencing symptoms in PTSD is the temporal dynamics model of emotional memory processing, set forth by Diamond et al. (2007). This model proposes that stressful experiences produce an intense, but brief, activation of memory-encoding plasticity within the hippocampus which is followed by a refractory period in which the structure may be thought of as having limited response potentialities. This refractory period isolates the memory by not producing a coherent narrative of the event but only a sharply focused memory of the trauma or its precursors. In conjunction with activation of the amygdala during the traumatic event, the implicit, fragmented, and primarily sensory structure of traumatic memories arise (Diamond et al., 2007).

Other interpretive models for intrusive memories in PTSD propose a more general information processing impairment underlying the intrusive memories characterizing PTSD. For instance, some have argued that people who suffer from PTSD have better visual imagery abilities which predispose them to experience more vivid traumatic imagery following a traumatic event (Bryant & Harvey, 1996; Stutmant & Bliss, 1985).

Although there seems to be a general acceptance of the fragmentary sensory phenomenology of traumatic memories, neural indications for these clinical reports during trauma recollection are scarce. If indeed traumatic memories consist of a vivid visual imagery of the event, neuroimaging studies should expect to find activation of neural regions underlying mental imagery, such as activation of the primary visual areas (Kosslyn et al., 1999; Cui et al., 2007). Functional neuroimaging studies conducted on PTSD patients when recalling the traumatic events have generally supported the hypothesis that emotional processing regions such as amygdala and insula are hyper-responsive in PTSD, while rostral and ventral portions of the medial prefrontal cortex (mPFC) are hypo-responsive (see Hughes & Shin, 2011 for a review). These findings were taken to reflect the exaggerated fear response and hyperarousal that patients experience when imagining the traumatic event. Among these studies, only few reported increased activity of visual areas in PTSD, which were proposed to underlie re-experiencing symptoms (Rauch et al., 1996; Lindauer et al., 2004). The abovementioned studies used methods characterized in low temporal resolution (including PET, SPECT and fMRI). Related research has also used magnetoencephalography (MEG), characterized in high temporal resolution that enables examination of the brain responses on the order of milliseconds. However, MEG studies have employed tasks that do not directly target patients’ traumatic memories. For example, a resting state paradigm showed elevation of delta activity over left temporal and right frontal regions in PTSD (Kolassa et al., 2007). Enhanced delta activity in the insula in PTSD was taken to reflect patients’ difficulty to identify and regulate their emotional states in response to trauma reminders. In addition, a series of experiments investigated the brain responses of PTSD sufferers to aversive pictures or words. They found facilitated sensory processing of affective stimuli between 170–210 ms in posterior regions in PTSD, indicative of a hypersensitive alarm system that is tuned for detection of potentially threatening cues in the environment (Rockstroh & Elbert, 2010). To our knowledge only one pilot study was conducted using MEG during recall of the traumatic memory. That study found decreased delta power in the secondary visual cortex, decreased beta power in the insula and decreased alpha power in the insula, premotor cortex and Broca’s area during trauma imagery, relative to a rest condition in PTSD. However, the generalizability of these findings is limited because the study included only nine women, all suffering from civilian PTSD, and no control group (Cottraux et al., 2015).

The present study aims at elucidating the neural oscillatory activity underlying trauma recollection using MEG in a paradigm resembling exposure therapy. We believe the sensitivity of MEG to a large spectrum of fast, oscillatory brain signals, combined with its high ability to map their anatomical origins, can contribute to a deeper comprehension of the nature of trauma recollection and its implementing neural mechanisms. We compared brain activity elicited by trauma recollection with a scripted narrative with that elicited by self-imagery in order to tap neural mechanisms responding to directed and internal trauma provocations. In addition, in order to investigate whether the brain mechanisms are specific to trauma provocation or constitute a more general information processing network, the oscillatory activity during exposure to a neutral event in PTSD patients was also be assessed. We hypothesized that exposure to trauma provocations will result in higher activity in perception-related brain regions, predominantly visual areas, that reflect the sensory aspect of traumatic memories in PTSD patients. Following previous functional neuroimaging studies, we further hypothesized greater activation in brain areas implicated in emotion and fear response such as the insula, and decreased activation in medial prefrontal cortical areas involved in modulation of fear responsiveness. Since the alpha rhythm is known to increase under conditions of mental relaxation (Klimesch, 1996) and to be suppressed during mental effort and in clinical symptoms of hyperarousal (Veltmeyer et al., 2006)), we predicted decreased alpha activity during traumatic compared to neutral conditions in both PTSD and healthy trauma-exposed controls, with higher alpha suppression in the PTSD group.

## Methods and Materials

### Participants

Seventeen PTSD patients and sixteen healthy trauma-exposed controls participated in the study. During the same MEG session, participants also underwent an oddball paradigm experiment (Herz et al., 2016). Two PTSD and one control participants dropped out during the task due to difficulty in withstanding test conditions. Additionally, three participants from the PTSD group and one from the control group were excluded from analysis due to excessive magnetic or muscle artifact. Hence, MEG analysis was based on 14 control participants (12 men, 2 women) and 12 PTSD participants (11 males, 1 woman). Exclusion criteria for both groups included symptoms or signs of psychosis or suicidality; drug/alcohol abuse in the previous 6 months; past history of brain injury, loss of consciousness or other neurological disease; and a contraindication to undergoing MRI or MEG. Groups were matched for age and time since the occurrence of the trauma. Participants experienced diverse adult trauma, including motor vehicle accidents (PTSD n=3, control n=7), terrorist attack (PTSD n=1, control n=1) and military related trauma (PTSD n=8, control n=6). Comorbidity of anxiety and depression was allowed in the PTSD group, given that diagnosis of PTSD preceded the comorbid diagnosis.

PTSD patients were recruited from the Outpatient Psychiatry Service of Hadassah Medical Center. Healthy control subjects were recruited from hospital staff and through advertisements in local media. Control group participants did not meet criteria for any psychiatric disorder and were not taking psychotropic medication at the time of the study. Comorbidity of anxiety and depression was allowed in the PTSD group, given that diagnosis of PTSD preceded the comorbid diagnosis. Antidepressant and anxiolytic/hypnotic medication were also allowed with the rationale that excluding medicated subjects would result in recruitment of a non-representative sample of patients. All participants were right handed. Time elapsed from the trauma was greater than one year. Participants were in good physical health confirmed by a complete physical and laboratory evaluation (including physical and neurological examination, vital signs, ECG, standard laboratory assays of blood and urine).

PTSD diagnosis was conferred using the Structural Clinical Interview for DSM-IV (SCID) (First et al., 1996), as the DSM-V was not published at the time of the study. All participants were further assessed using the DSM-IV Clinician Administered PTSD Scale (CAPS) (Blake et al., 1995), Hamilton Depression Rating Scale (Hamilton 1960), and Hamilton Anxiety Rating Scale (Hamilton, 1959). All procedures were in compliance with the Code of Ethics of the World Medical Association (Declaration of Helsinki) and approved by the Hadassah Hebrew University Medical Center Ethics Committee. Written informed consent was obtained from all subjects.

### Procedure

Individualized trauma scripts and a common neutral script, approximately 30 seconds long each, were devised according to established procedure (Pitman, 1987). Individualized trauma scripts portraying actual experiences from each participant’s past were composed in advance based on the participant’s description of the event. They were written in second person, singular, present tense, incorporating bodily responses, cognitions and emotions experienced during the traumatic event.

Scripts were recorded by the same neutral male voice for playback during the MEG scan. The traumatic and neutral script-imagery conditions were repeated twice. Participants lay supine with their eyes closed during the entire experiment in order to minimize eye movement artifacts. A baseline resting period (1 min) was followed by four experimental blocks. Each block proceeded as follows: 1. Narrative script: participants listened through headphones to the traumatic or neutral script (30 sec), 2. Imagery: participants were encouraged to form a vivid mental image of the script previously heard, incorporating auditory, somatosensory, olfactory or visual sensations that were associated with the event (30 sec), 3. Rest: participants lay still and let go of the remembered event. Figure 1 outlines the flow of the experimental session.

**Figure 1.**
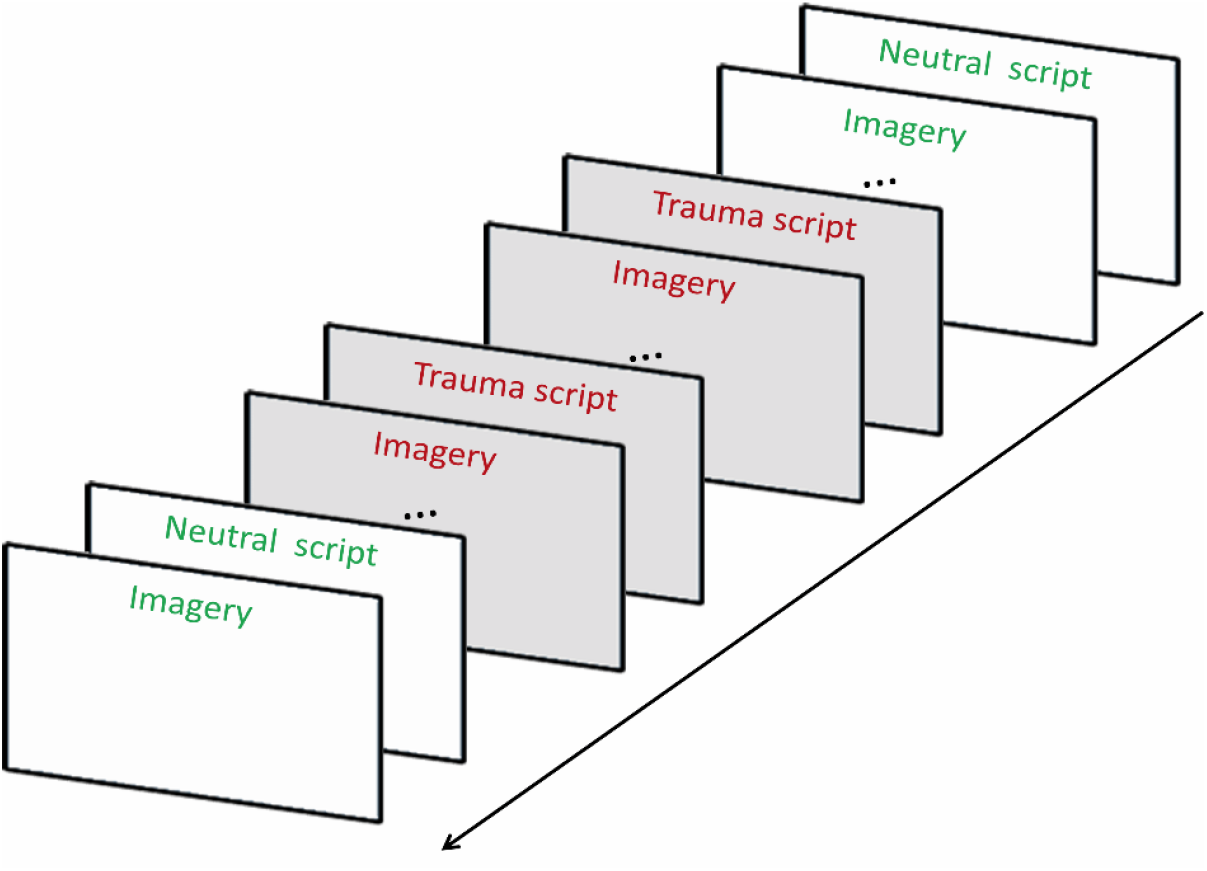
Flow of events in the script-driven imagery paradigm. Each block started and ended with a 60-seconds resting period, after which a neutral or individualized trauma script was presented. Reading of the scripts lasted ~30 seconds. Immediately after presenting the script an imagery condition started, in which the participant was encouraged to remember olfactory, auditory, somatosensory, and visual sensations that were associated with the event.

### MEG Data Acquisition

MEG recordings were conducted using a whole-head, 248-channel magnetometer array (4-D Neuroimaging, Magnes 3600 WH) in a magnetically shielded room. Reference coils located a short distance (~30 cm) away from the 248 channels, and oriented by the x, y and z axes, were used to remove environmental noise. Prior to data acquisition, three localization coils were placed at the two preauricular points and nasion to localize the participant’s head relative to the MEG sensors. Head-shapes were digitized using a Polhemus Fastrack digitizer. The data were digitized at a sample rate of 1017.25 Hz and a 1-400 Hz online band-pass filter was used. An additional channel recorded the 50 Hz signal from the power outlet which was used to clean the mains noise and its harmonics by calculating the average 50 Hz cycle on every MEG channel and removing it from the data, hence allowing the cleaning of the line power noise without a notch-filter (Tal & Abeles, 2013).

### MEG analysis

Two malfunctioning sensors were discarded from all analysis. Heartbeat artifacts were removed using an event-synchronous cancellation algorithm, implemented by Tal and Abeles (2013). Each epoch was visually examined for muscle and jump (in the MEG sensors) artifacts. Trials containing power jumps were discarded. Muscle artifacts were manually rejected by applying a high-pass filter of 60Hz and identifying deviant trials based on their variance. Spatial Independent Component Analysis (ICA) (Jung et al, 2000) was applied in order to identify eye movements and blink artifacts. Trials containing these artifacts were rejected from the data.

Data were analyzed with MATLAB (The Mathworks, Andover, MA) using Fieldtrip toolbox (20). Data were separated into the different conditions (four ‘narrative script’ and four ‘imagery’ conditions). Each condition was segmented into 2000 ms epochs, with 1500 ms overlap (in 500ms steps). For source estimation, individual MRIs were fitted to the digitized head shapes using AFNI (21). Synthetic aperture magnetometry (SAM) beamformer (Robinson & Vrba, 1999) was applied at six frequency bands: delta (1–4 Hz), theta (4–7 Hz), alpha (8–12 Hz), beta (13–30 Hz), gamma (30-80 Hz) and high gamma (80-150), based on the covariance matrix of all segments. SAM is a nonlinear minimum-variance beamformer algorithm. It uses the signal covariance calculated from the magnetic signals recorded at MEG sensors to construct optimum spatial filters at each voxel in the brain, by minimizing the correlations with all other analyzed voxels. Estimation of the source power for each voxel results in a 3D spatial distribution of the power of the neuronal sources.

For each participant in our experiment we calculated a global covariance matrix adopting the segmented data of 2000ms for each frequency band of interest. The beamformer weights were calculated based on the above covariance matrix using data from all trials. Then for each frequency band, weights were multiplied by the data, thus creating “virtual sensor” time-series for every condition. The data was normalized by dividing it by a noise estimate, which consisted of the variance of all trials. In order to facilitate group analysis, head models were constructed by co-registering each participant’s SAM volume to his previously obtained MRI scan based on the position of the fiduciary markers established during the digitization phase. Each participant’s MRI image and its co-registered SAM volume were then transposed into a common anatomical space (Talairach).

In order to test for differences between groups, voxel-level group statistics was conducted using AFNI (Cox, 1996). Results were corrected for multiple comparisons based on the Random Field theory methods (Pantazis et al., 2005). A simulation function (ClustSim), determining the probability to get significant clusters at random given a template brain and specific spatial resolution was used. Following previous studies suggesting smoothing of 5 mm to be the most accurate parameter for MEG data (Pantazis et al, 2005; Barnes et al., 2004) we used a spatial smoothing of 7 mm in the simulation for even more conservative results. According to this simulation, given clusters of voxels with a p-value smaller than 0.05 and alpha significance level of p < .01, clusters exceeding 41 voxels do not count as random noise. The voxels averaged activity in clusters exceeding that cluster size was used in the following analyses.

Clusters with significant effects were identified by a voxel-level F-test with valence (traumatic, neutral) and sequence (first, second presentation) as the within-subject factors, and group (PTSD, control) as a between-subjects factor. The power at each frequency band was used as the dependent variable. Separate analyses were performed for the narrative listening (‘script’) and the imagery segments.

## Results

### Descriptive measures

Demographic information and psychological assessment results can be found in Table 1. The control and PTSD groups did not differ on age (t(24) = 0.79, n.s) or time since the occurrence of the trauma (t(24) = 1.85, n.s). PTSD symptoms scores of patients were in the severe range while control participants were not symptomatic. Clinical ratings showed, as anticipated, that PTSD patients had significantly higher PTSD (t(24) = 11.05, p < .001), depression (t(24) = 5.35, p < .001) and anxiety symptoms (t(24) = 7.98, p < .001) compared with controls.

**Table 1.**
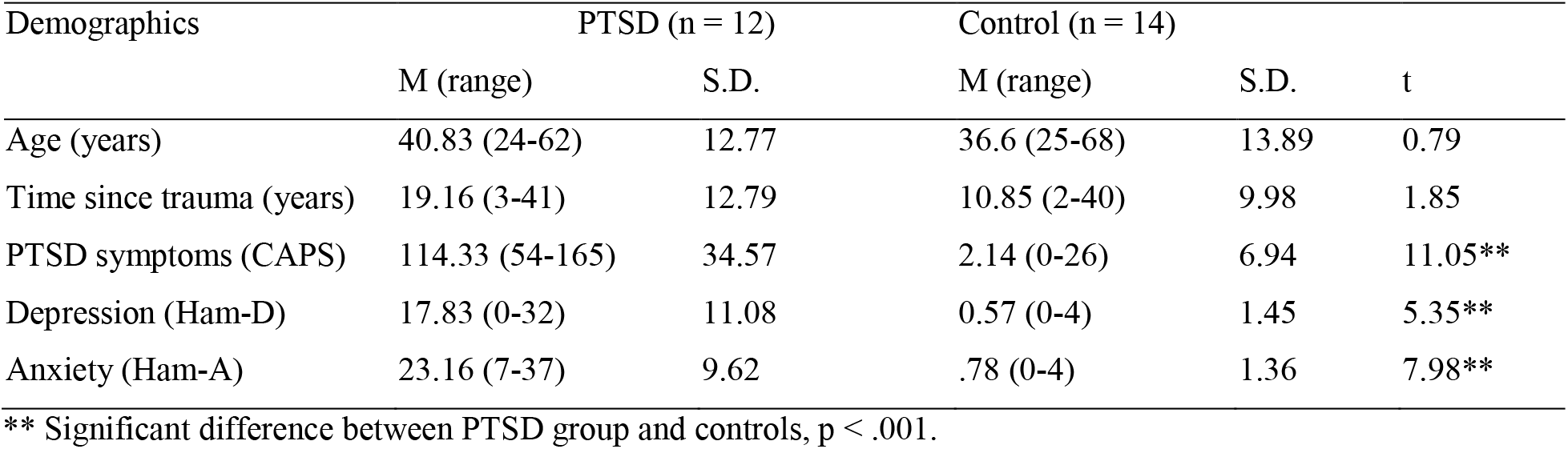
Age, Time since Trauma and Symptom Reports

### MEG Results

#### Group main effect

Across stimuli valence (neutral and traumatic conditions), PTSD participants showed less gamma and high-gamma activity compared to controls. Regions with decreased activity were similar for narrative and imagery, and included visual areas, as well as frontal and post-central regions (in Table 2).

**Table 2.**
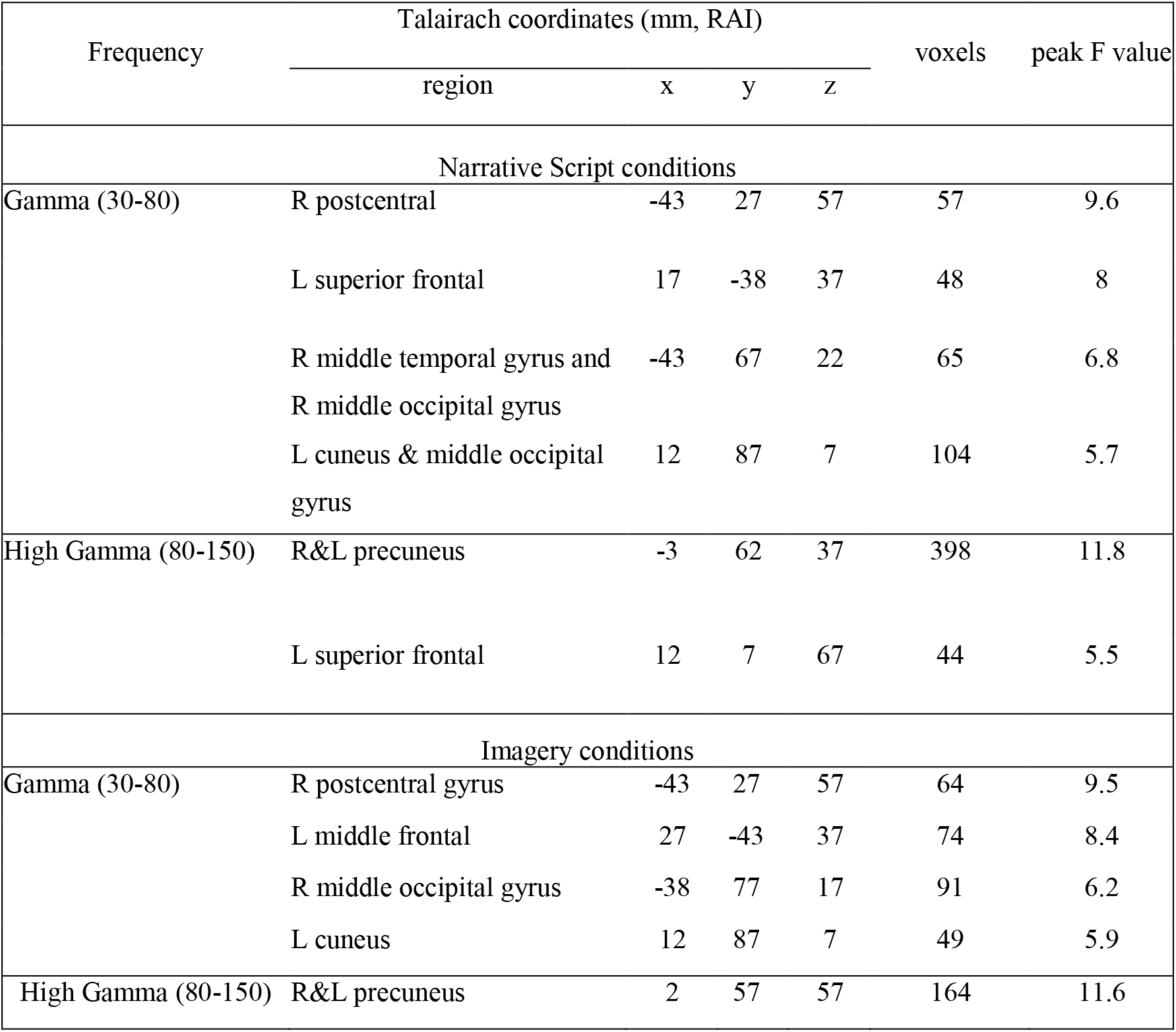
Significant clusters showing group main effects

#### Group X Valence interaction effect

Overall, a consistent pattern of group X valence interaction emerged: PTSD participants exhibited higher power during the traumatic condition (both during narrative and imagery), while the control group showed equivalent or less power during the traumatic compared to the neutral condition. This pattern of interaction was found across all frequency bands. The detailed results relating to this group X valence interaction will be described in turn for each frequency band separately and can be seen in Tables 3–4. Effects not reflecting group differences are reported in Supplementary Material 1.

**Table 3.** Group X Valence Interaction Effects in Narrative Script conditions

| Frequency | Talairach coordinates (mm, RAI) |  |  |  | voxels | peak F value |
| --- | --- | --- | --- | --- | --- | --- |
|  | region | x | y | z |  |  |
| Delta (1-4) | R superior, middle and medial frontal | -8 | -68 | 17 | 206 | 15.1 |
|  | L inferior and superior temporal | 62 | 7 | -18 | 189 | 13.3 |
|  | R. middle and superior temporal | -53 | 22 | 2 | 130 | 7.2 |
| Theta (4-7) | R medial frontal and anterior cingulate | -3 | -23 | -18 | 49 | 16 |
|  | R medial, superior and middle frontal | -23 | -48 | 12 | 67 | 14.1 |
|  | L medial and superior frontal | 12 | -48 | 12 | 56 | 11.8 |
|  | R middle and superior temporal | -58 | 22 | -3 | 98 | 9.3 |
|  | L middle and superior temporal | 52 | -3 | -18 | 236 | 9.1 |
|  | L cingulate gyrus | 12 | 7 | 37 | 79 | 8.3 |
| Alpha (8-12) | R superior and medial frontal | -8 | -68 | 17 | 200 | 11.5 |
| Beta (15-30) | L inferior temporal and fusiform gyrus | 57 | 57 | -13 | 120 | 11 |
|  | R middle and inferior frontal gyrus | -38 | -53 | -8 | 258 | 10.7 |
| Gamma (30-80) | L inferior and middle occipital | 42 | 82 | -13 | 55 | 6.7 |
|  | L uncus and inferior temporal | 27 | 7 | -38 | 41 | 4.8 |
| High Gamma (80-150) | R&L cuneus | 12 | 87 | 37 | 647 | 9.7 |
|  | L parahippocampal, L middle and inferior temporal | 42 | 32 | -8 | 199 | 4.7 |

**Table 4.** Group X Valence Interaction Effects in Imagery conditions

| Frequency | Talairach coordinates (mm, RAI) |  |  |  | voxels | peak F value |
| --- | --- | --- | --- | --- | --- | --- |
|  | region | x | y | z |  |  |
| Delta (1-4) | L cuneus | 2 | 92 | 7 | 64 | 13.3 |
|  | R cingulate gyrus | -8 | -28 | 27 | 77 | 9.5 |
|  | L postcentral gyrus | 37 | 27 | 57 | 48 | 7 |
| Theta (4-7) | R superior frontal, medial frontal and cingulate gyrus | -13 | -43 | 32 | 513 | 13.4 |
|  | L middle temporal | 57 | 32 | -13 | 79 | 9.3 |
|  | R middle temporal gyrus | -63 | 2 | -8 | 41 | 7.4 |
| Alpha (8-12) | R anterior cingulate | -18 | -38 | -3 | 96 | 7.9 |
| Beta (15-30) | L superior frontal | -22 | 38 | 27 | 166 | 14.4 |
|  | L middle temporal and parahippocampal gyrus | 27 | 27 | 2 | 255 | 9.2 |
|  | R insula | -28 | -18 | 12 | 58 | 8.1 |
|  | R thalamus, parahippocampal and fusiform gyrus | -28 | 27 | 2 | 91 | 6.6 |
| High Gamma (80-150) | R&L cuneus | 12 | 77 | 22 | 1002 | 8.5 |

##### High-gamma band (80-150 Hz)

In the high-gamma band, PTSD participants exhibited widespread abnormal recruitment of the occipital regions, peaking at the cuneus. In this area, high-gamma power was larger during the traumatic script listening and imagery compared to the neutral conditions in the PTSD group, whereas the control group had equivalent high-gamma power during the trauma and neutral script listening and imagery (Fig. 2).

**Figure 2.**
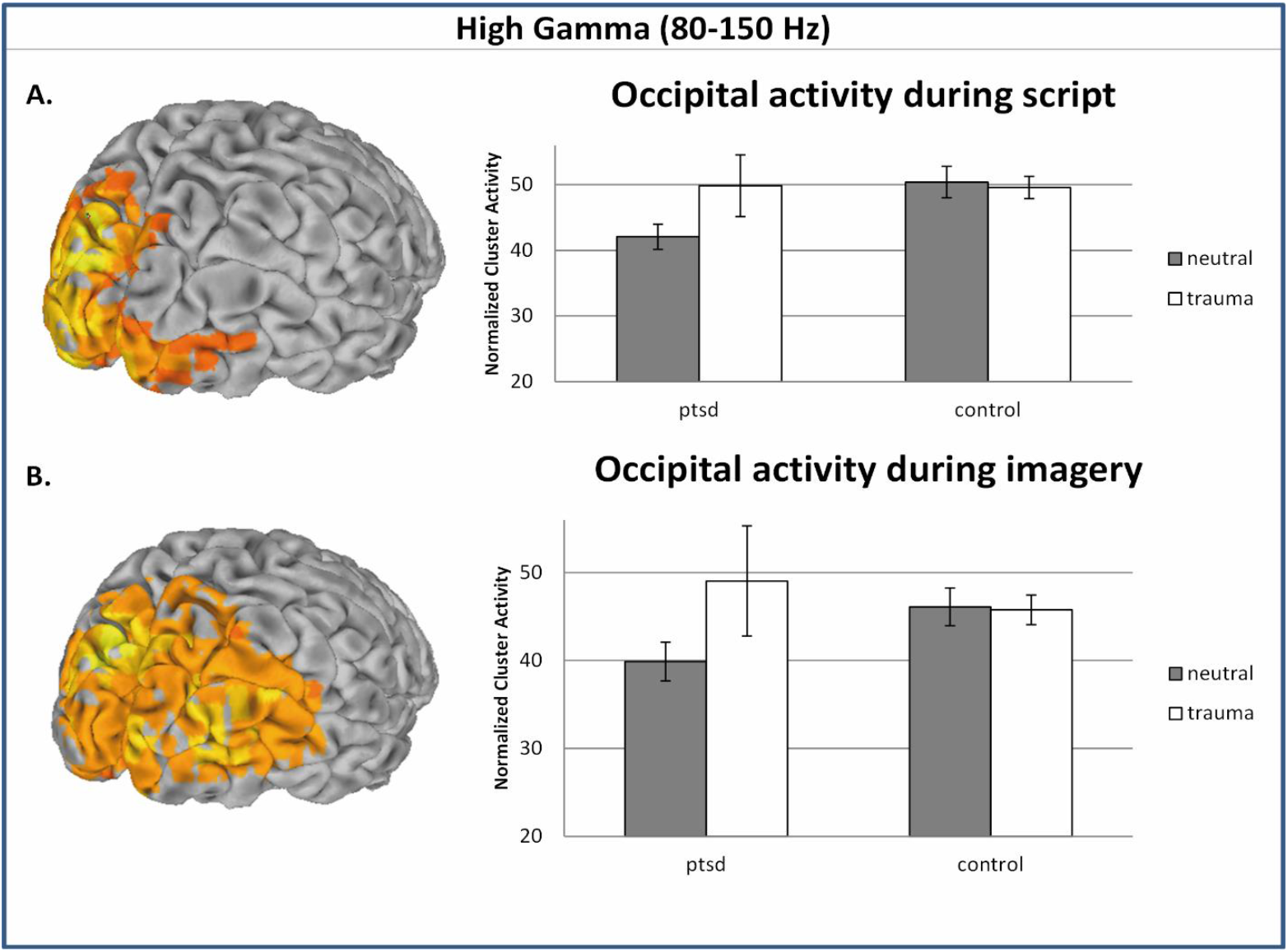
Altered high-gamma modulation of PTSD participants in visual areas peaking at the cuneus. Colored areas indicate clusters with significant interacion effects. Bar graphs depict mean cluster activity elicited by neutral (grey) and trauma (white) narratives during listening (A) and imagining (B).

##### Gamma band (30-80 Hz)

In the gamma band, the same pattern of group X valence interaction was found in the left inferior and middle occipital gyri and in the left inferior temporal gyrus; PTSD patients had higher gamma power in response to trauma script compared to neutral script, whereas the controls did not show such differential power response.

##### Beta band (13-30 Hz)

The same pattern of group X valence interaction was found in the beta band, over limbic and frontal regions. During imagery, PTSD patients had higher beta power in response to trauma imagery compared to neutral imagery over the right and left parahippocampal area (Fig. 3), right insula and left superior frontal gyrus, while the control group did not show such differential power response. During the narrative, PTSD patients had higher beta power in response to the trauma script compared to the neutral one over the right middle and inferior frontal gyrus and left inferior temporal gyrus, whereas the control group did not show such differential power response.

**Figure 3.**
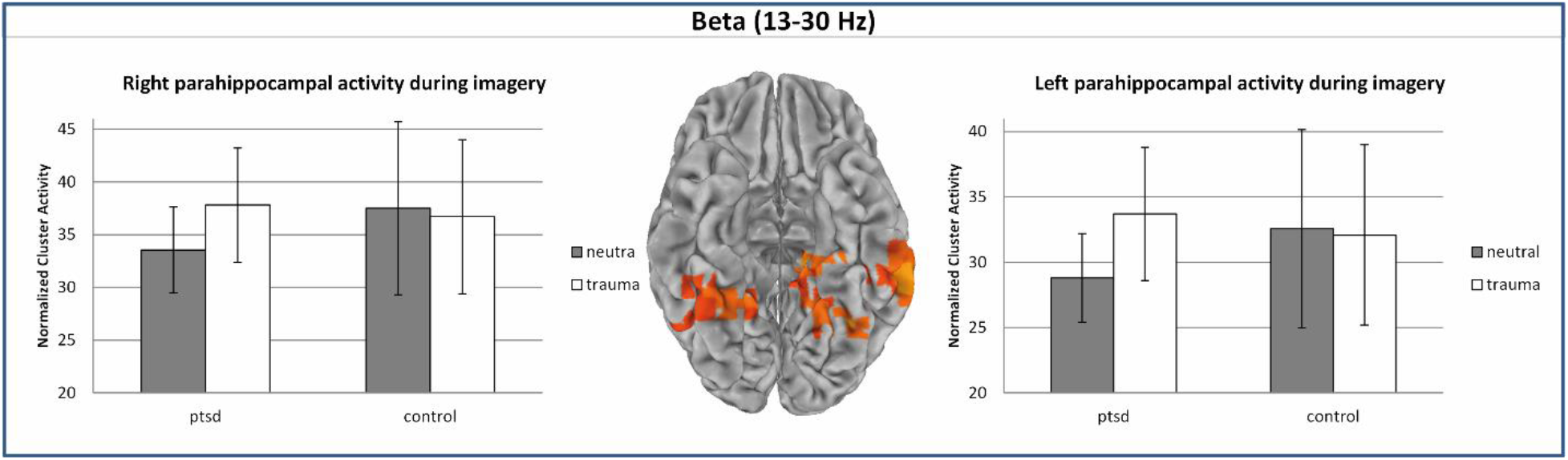
Altered modulation of the beta activity in PTSD patients in the right and left parahippocampal regions during script imagination. Colored areas indicate clusters with significant interacion effects. Bar graphs depict mean cluster activity elicited by neutral (grey) and trauma (white) narratives.

##### Alpha band (8-12 Hz)

In the control group, alpha suppression during the traumatic conditions (both narrative and imagery) relative to the neutral conditions was observed in frontal regions. In the PTSD group, however, alpha power was higher in the traumatic conditions (both narrative and imagery) compared to the neutral conditions (Fig. 4).

**Figure 4.**
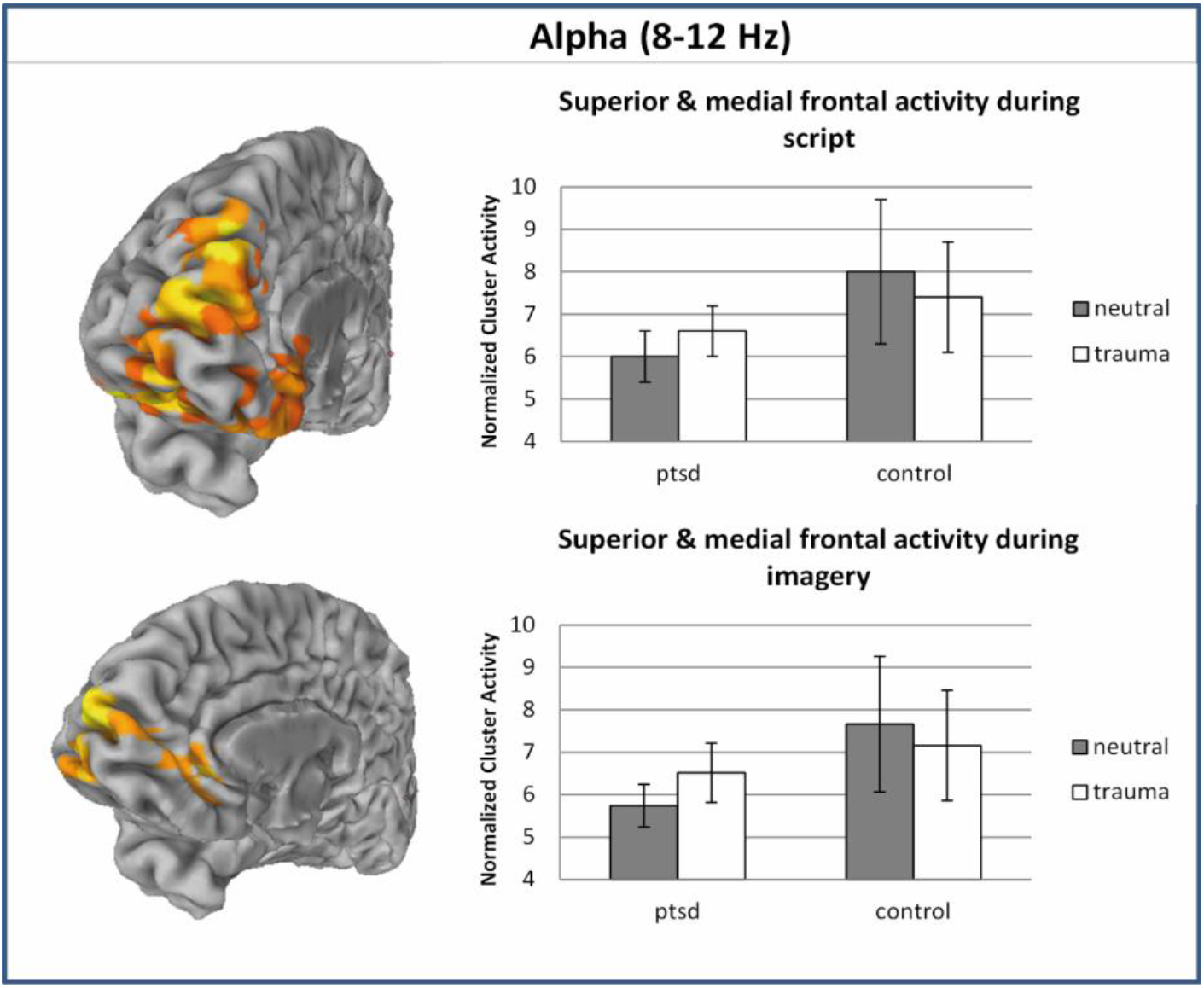
Altered modulation of the alpha activity in PTSD participants compared to controls in medial frontal regions. Colored areas indicate clusters with significant interacion effects. Bar graphs depict mean cluster activity elicited by neutral (grey) and trauma (white) narratives during listening (upper) and imagining (lower).

##### Theta band (4-7 Hz)

The same pattern of interaction was observed in the theta band, over frontal and temporal regions. During the imagery condition, PTSD patients had higher theta power during the trauma imagery compared to neutral imagery in the medial frontal region and cingulate gyrus and in bilateral middle temporal gyrus, whereas controls showed the opposite trend. During the narrative, PTSD patients had higher theta power for the trauma script compared to neutral script in the superior and medial frontal regions including the cingulate gyrus and in bilateral superior temporal gyrus, whereas controls exhibited the opposite pattern.

##### Delta band (1-4 Hz)

In response to script, PTSD patients had higher delta power in response to trauma script compared to neutral script over the bilateral superior temporal and prefrontal cortex, while the control group did not show such differential power response.

In response to imagery, the same pattern of interaction was evident in the right cingulate and left cuneus. However, in the postcentral gyrus the opposite pattern was found, with PTSD patients having less delta power in response to trauma compared to neutral imagery, whereas controls had equivalent power across trauma and neutral imagery.

## Discussion

The results of the present study provide evidence for a distinct neural processing of traumatic memories among individuals suffering from PTSD compared to healthy trauma-exposed controls. It was found that PTSD patients show increased power in high-frequency bands (gamma and high-gamma) across the visual regions, peaking at the primary visual area, when listening to and imagining their trauma narrative compared to a neutral event. In comparison, control participants had equal gamma and high-gamma power when recalling their traumatic experience and a neutral event (Fig. 2). The gamma rhythm has been suggested to underlie the mutual information transfer between brain regions (Başar-Eroglu et al., 1996), and to have a critical role in the binding of stimulus information for coherent perception (Lutzenberger et al., 1995). Increased gamma and high-gamma power of the PTSD group over visual areas may reflect the vivid mental images usually described by patients when recalling their traumatic event (Ehlers et al., 2002), and point to the possible role of occipital regions in the pathophysiology of PTSD that may be overlooked when using neuroimaging techniques characterized by low temporal resolution. Previous MEG studies have also found altered activity in visual processing areas in traumatized individuals in response to aversive, not trauma related, pictures. One study found hypoactivity of occipital areas in survivors of war and torture, that was suggested to represent a defensive reaction and adaptive adjustment of the brain to exposure for a threatening environment (Catani et al., 2009). A dampened posterior activity was also suggested to reflect pathological inhibition of emotional visual processing, triggered by hyper-reactivity to potentially threatening stimuli (Rockstroh & Elbert, 2010). While the current study incorporated direct exposure to the personalized traumatic memory and not to general aversive stimuli, the results of also point toward decreased activation (power) of visual regions in PTSD in response to neutral recollection compared to controls (Fig. 2). Interestingly, we also found PTSD patients to exhibit less high-gamma activity across stimuli valence (i.e. group main effect) both during narrative and imagery in the precuneus, a region proposed to facilitate the transfer of sensory-bound representations into more elaborated contextualized representations that can be integrated with existing autobiographical information (Brewin et al., 2010).

Apart from altered activity over visual areas, we also found abnormal recruitment of superior temporal areas that overlap with the auditory cortex in the delta and theta bands in PTSD. PTSD patients had higher delta and theta power during trauma imagery or trauma script listening compared to neutral recollection, while in controls no difference between trauma and neutral recollection was present. The convergence of findings over perceptual brain areas is in agreement with recent resting-state MEG study in PTSD (Badura-Brack et al., 2017), and might index altered perceptual experience of PTSD patients when recalling their traumatic event. In addition, we found abnormal modulation of the lower frequencies over frontal regions in PTSD. PTSD patients had increased alpha and theta power mostly over the medial prefrontal cortex (mPFC) in response to trauma narrative and imagery compared to neutral recollection. Controls exhibited the opposite trend, with theta and alpha suppression in response to trauma imagery or trauma script listening compared to neutral recollection (Fig. 4). The alpha suppression in response to traumatic stimuli found in the control group is in accordance with our expectations, as alpha was previously shown to be suppressed during mental effort and hyperarousal (Klimesch, 1996; Veltmeyer et al., 2006). In the PTSD group, however, alpha power increased in response to the traumatic conditions but was still continuously suppressed compared to controls (Fig. 4). The presence of this general suppression of frontal activity in PTSD fits well with the literature, pointing to dysregulation of the mPFC as a key factor in the generation of PTSD symptoms. Current models propose that inadequate top-down control by the mPFC over limbic regions cause perpetuation of hyperarousal and reactivity in PTSD (Liberzon & Sripada, 2007). In agreement with this model, most of the symptom provocation studies have implicated hyperactivation of paralimbic and limbic structures in the symptomatic state of PTSD. In our study, some paralimbic areas exhibited altered activity in the PTSD group, primarily in the beta band. These areas included the bilateral parahippocampal gyrus and right insula, in which decreased beta power during neutral imagery but increased beta power during trauma imagery was found in PTSD (Fig 3).

Interestingly, the abnormal brain activity of PTSD patients compared to controls was remarkably similar during script-listening and script-imagery conditions. The recruitment of perceptual regions, both when patients are exposed to explicit cues of the trauma, as well as when asked to form a mental imagery of the event, suggests that mental imagery leads to comparable intensity of memory recollection as in response to external triggers. Indeed, studies comparing imaginal versus in-vivo exposure in patients suffering from specific phobia found that imaginal exposure was just as effective as traditional in-vivo exposure therapy (Hunt & Fenton, 2007). These findings suggest that exposure therapies in PTSD may be beneficial by incorporating only one exposure modality (either in-vivo or imaginal exposure). Also, in a study assessing the characteristics of intrusive memories and their change over the course of therapy, it has been shown that the vividness of the traumatic memories faded gradually (Hackman et al, 2004). If indeed activation of perceptual brain regions corresponds with the vividness of the traumatic memory, such improvement may be indexed by lower recruitment of perceptual brain regions during therapy and to ultimately reach equivalent activation during recall of the trauma as when recalling neutral events, in a similar manner to the brain activation characterizing healthy trauma-exposed controls.

In conclusion, the current study indicates that the neural processing of PTSD patients when recalling their traumatic memory is different than the one underlying recollection of stressful events in healthy individuals. Altered modulation of high-frequencies over visual areas in PTSD might reflect the vivid mental imagery taking part during the trauma recollection, while altered modulation of the lower frequencies over medial frontal areas may reflect impaired top-down regulation over limbic regions. Further studies are needed in order to replicate these results in a larger sample size, taking into account the subjective reports of patients while recalling their traumatic event. Investigating the fast oscillatory neural dynamics of PTSD patients during trauma recollection can help us better understand how they re-experience their trauma through memory, and may aid in monitoring the efficacy of psychotherapy interventions incorporating exposure components in their procedure and in developing new therapeutic interventions to decrease the vividness of the traumatic memories.

## Acknowledgments

This work is partly based on a PhD thesis submitted by NH to the Department of Psychology, Bar-Ilan University, under supervision of AG and TP.

