## Supplemental information for "Neural oscillations while remembering traumatic memories in PTSD"

### Supplemental Results

While the main interest in our study was the differential brain responses of PTSD patients compared to control, we portray here other effect that may also be of interest.

**Script effect:**

Differences between traumatic and neutral conditions across groups can identify brain regions involved in trauma processing in general. We found less power in the lower-frequency bands in response to traumatic conditions compared to neutral conditions in frontal areas (theta, alpha, beta) and posterior and visual areas (delta, theta, and beta). However, in the higher-frequency bands of gamma and high-gamma, increased power was found in response to the traumatic conditions, mostly over the superior frontal and orbitofrontal gyri (Fig. S1). Regions in which differences between neutral and traumatic conditions were found can be seen in Tables S2 and S3.

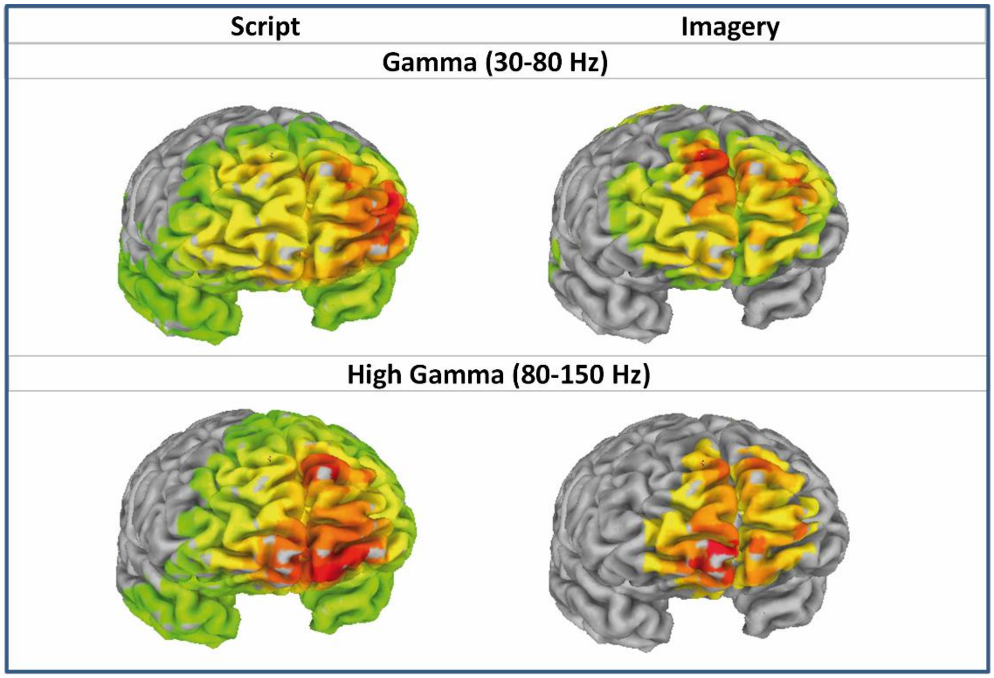


**Figure S1.** Increased activation in the gamma and high-gamma frequencies over frontal and orbitofrontal regions in response to traumatic compared to neutral conditions (across groups).

**Table S2. Script Main Effects in Narrative Script conditions**

| Frequency | Talairach coordinates (mm, RAI) | | | | voxels | peak F value |
|---|---|---|---|---|---|---|
| | region | x | y | z | | |
| Delta (1-4) | R lingual, R&L cuneus | -3 | 72 | 2 | 1372 | 19.7 |
| Theta (4-7) | R&L cuneus | 7 | 92 | 12 | 1275 | 19.4 |
| | R&L superior frontal | 12 | -23 | 57 | 313 | 15.5 |
| Alpha (8-12) | L cingulate and R superior frontal | 22 | 2 | 27 | 872 | 18.5 |
| Beta (15-30) | L superior occipital gyrus and L cuneus | 42 | 77 | 27 | 175 | 10.9 |
| | R&L superior frontal | 22 | -13 | 57 | 157 | 14.3 |
| Gamma (30-80) | R&L middle and medial frontal gyrus, L&R middle and superior temporal. | -26 | -57 | 12 | 5296 | 26.2 |
| High Gamma (80-150) | L superior frontal | 7 | -63 | -13 | 6090 | 21.4 |

**Table S3. Script Main Effects in Script Imagery conditions**

| Frequency | Talairach coordinates (mm, RAI) | | | | voxels | peak F value |
|---|---|---|---|---|---|---|
| | region | x | y | z | | |
| Delta (1-4) | R&L Postcentral gyrus | 17 | 42 | 72 | 1908 | 31.5 |
| Theta (4-7) | L precuneus | 7 | 62 | 27 | 2735 | 27.29 |
| Alpha (8-12) | R&L precentral gyrus | 27 | 12 | 67 | 981 | 15.5 |
| Beta (15-30) | R&L medial frontal gyrus, cingulate, middle frontal, and superior frontal | -3 | -23 | 47 | 2417 | 35.9 |
| Gamma (30-80) | R&L superior and medial frontal | -13 | -58 | 27 | 1810 | 16.9 |
| | R&L postcentral and precentral gyrus | 22 | 42 | 67 | 536 | 17.8 |
| High Gamma (80-150) | R middle temporal & L cingulate | -33 | 57 | 17 | 715 | 6.6 |
| | R&L superior and medial frontal | -13 | -68 | -3 | 365 | 8.7 |
| | L middle occipital | 47 | 72 | -3 | 275 | 7.2 |

### Group X Script X Sequence effect:

In order to test whether the order of presentation had any effect on the pattern of Group X Script interactions, we examined the Group X Script interaction for the first vs. second presentation (Group X Sequence X Script interaction). Although in several frequency bands the order had some influence on the intensity of the differences between conditions, it did not change their pattern. For example, while in the alpha band lower alpha activity of PTSD patients compared to controls was found mainly for the neutral conditions (Group X Script interaction, Fig. 5 in the main text), the additions of the sequence effect revealed that these differences were more pronounced in response to the first neutral presentation than the second. Since the order of presentation did not change the direction of the effects (i.e. ordinal rather than disordinal interaction), it was not treated as confounding factor. Regions in which Group X Sequence X Script interactions were found can be seen in Tables S4 and S5.

**Table S4. Group X Script X Sequence Interaction Effects in Narrative Script conditions**

| Frequency | Talairach coordinates (mm, RAI) | | | | voxels | peak F value |
|---|---|---|---|---|---|---|
| | region | x | y | z | | |
| Delta (1-4) | R supramarginal and R inferior parietal gyrus | -53 | 57 | 32 | 73 | 13.1 |
| Theta (4-7) | R&L cingulate gyrus and R. medial frontal gyrus | -3 | -18 | 42 | 170 | 11.7 |
| | R&L precuneus | 2 | 67 | 52 | 116 | 13.1 |
| | R inferior parietal lobule | -43 | 47 | 42 | 66 | 8.7 |
| | R inferior & middle occipital, R lingual gyrus | -28 | 87 | -3 | 57 | 16.5 |
| | L inferior parietal lobule | 52 | 32 | 37 | 42 | 11.3 |
| Alpha (8-12) | L cingulate | 12 | 2 | 32 | 123 | 15.1 |
| | R inferior parietal lobule and R precuneus | -38 | 42 | 42 | 118 | 14.4 |
| | R&L precuneus | -3 | 72 | 47 | 69 | 11.2 |

**Table S5. Group X Script X Sequence Interaction Effects in Script Imagery conditions**

| Frequency | Talairach coordinates (mm, RAI) | | | | voxels | peak F value |
|---|---|---|---|---|---|---|
| | region | x | y | z | | |
| Theta (4-7) | L superior occipital gyrus | 32 | 77 | 32 | 106 | 16.2 |
| Beta (15-30) | R Insula | -33 | -13 | 12 | 48 | 7.9 |
| Gamma (30-80) | R middle and superior temporal, R insula | -43 | 32 | -3 | 751 | 8.5 |
| | R cingulate gyrus | -18 | 17 | 42 | 62 | 7.4 |
| High Gamma (80-150) | R postcentral, R superior and middle temporal | -18 | 32 | 62 | 761 | 7 |
